# White matter plasticity during second language learning within and across hemispheres

**DOI:** 10.1101/2023.04.21.537810

**Authors:** Xuehu Wei, Thomas C. Gunter, Helyne Adamson, Matthias Schwendemann, Angela D. Friederici, Tomás Goucha, Alfred Anwander

## Abstract

Adult second language (L2) learning is a challenging enterprise inducing neuroplastic changes in the human brain. However, it remains unclear how the structural language connectome and its subnetworks change during adult L2-learning. The current study investigated longitudinal changes in white matter (WM) language networks in each hemisphere, as well as their interconnection, in a large group of Arabic-speaking adults who learned German intensively for six months. We found a significant increase in WM-connectivity within bilateral temporal-parietal semantic and phonological subnetworks and right temporal-frontal pathways mainly in the second half of the learning period. At the same time, WM-connectivity between the two hemispheres decreased significantly. Crucially, these changes in WM-connectivity are correlated with L2 performance. The observed changes in subnetworks of the two hemispheres suggest a network reconfiguration due to lexical learning. The reduced interhemispheric connectivity may indicate a key role of the corpus callosum in L2-learning by reducing the inhibition of the language-dominant left hemisphere. Our study highlights the dynamic changes within and across hemispheres in adult language-related networks driven by L2 learning.

**Significance:** The neuroplastic changes induced by learning a second language (L2) in adulthood open up new perspectives for understanding brain function. The current study shows structural changes in the language network of Arabic native speakers who learned German intensively in two phases of three months each. We found a marked change in the left-hemispheric lexical-semantic language system and the right fronto-temporal pathway, accompanied by a decrease in white matter connectivity in the corpus callosum during L2 learning, which occurred mainly in the second period of L2 acquisition. The reduced interhemispheric connectivity suggests that the inhibitory role of the corpus callosum, relevant for native language processing, is reduced by L2 learning. Our findings demonstrate a clear experience-dependent structural plasticity in the human brain during L2 learning.

## Introduction

Cognitive functions develop in parallel with the plastic adaptation of the brain (1–6). This suggests that the gray and white matter of the brain is altered by the acquisition of new skills and thus is modulated by lifelong experiences, such as the acquired native language (7). Second language (L2) learning in adulthood is a complex task that requires the adaptation of multiple brain systems related to a wide range of novel tasks to be mastered. To date, changes associated with L2 learning were reported to extend beyond the brain regions of the native language network in the left-hemisphere (8–10), with additional involvement of the right-hemisphere (11, 12), as well as plasticity in the white matter connections between the two hemispheres (13, 14). How these changes in the gray and white matter might develop during L2 learning is described in a model (15) called the Dynamic Restructuring Model (DRM).

The DRM-model postulates three distinct phases of structural adaptation that depend on the quantity and quality of the language learning and language switching experience. In the earliest phase, L2 learning leads to changes in the gray matter areas that support the processing of the new language. Next, in the intermediate consolidation phase, the white matter pathways connecting the language processing areas show a structural modulation. Finally, in the peak efficiency phase, the model predicts further changes in brain structure, including increased frontal white matter connectivity, leading to highly efficient L2 processing and language switching performance. However, longitudinal studies of white matter changes in L2 learning in large samples of adults who have achieved proficiency beyond the beginner level are still lacking. The present study aims to investigate different phases of longitudinal white matter changes within each hemisphere and across hemispheres, to clarify the role of the corpus callosum, and to describe the brain mechanisms involved in L2 learning.

L2 learning comprises the acquisition of a new vocabulary, which includes learning novel phonemes, phonetic categories as well as word meanings, in addition to a new grammar. At the behavioral level, it has been previously reported that lexical-semantic processing of newly learned words and simple grammar is relatively easy to acquire, and native-like performance can be achieved in L2 learners (16). In contrast, it is more difficult for late L2 learners to perform real-time syntactic analysis, and they do not achieve automatic, highly proficient syntax processing until a late stage of learning (12, 16).

At the neurofunctional level, brain imaging studies have shown that low proficient L2 learners have less overlap in brain activation between first and second language processing than high proficient L2 learners (17) and recruit additional brain areas in the right hemisphere (18). These brain areas may support language proficiency by effectively handling word retrieval (11). Comparing first and second language brain activation in the lexical-semantic domain was found to depend on the learners’ performance, but differences in the grammatical domain on the age of L2 acquisition (19).

Studies focusing on the neuroplasticity of the language system as a function of L2 learning (9, 10) have reported changes in the gray matter of the bilateral Inferior Frontal Gyrus (IFG), Inferior Parietal Lobe (IPL), and anterior and posterior Temporal Lobe (TL) (13, 20–23). In particular, they include cortical gray matter changes in the bilateral TL and IPL, related to phonological and lexical-semantic memory systems that are crucial for the acquisition of the new vocabulary (24–28). Additionally, since languages differ in their syntactic and morphological rules, successful L2 acquisition also depends on the brain’s adaptation to grammatical processing (29, 30). In native language processing, semantics and grammatical rules are processed in a left-lateralized network including inferior frontal and temporal-parietal regions, which are connected via dorsal and ventral white matter pathways (31). L2 acquisition during adulthood requires neural adaptations that reach beyond the classical language network, involving the right hemisphere (8–10), playing an essential role in the early learning phases when L2 processing is not yet fully automatized (24, 32).

In addition to the reported changes in gray matter, plasticity of the white matter language pathways in L2 learning (9) has also been suggested in previous cross-sectional studies comparing bilinguals and monolinguals (20, 23, 33), as well as in some longitudinal language learning studies (13, 22). These studies have shown an association between L2 acquisition and local changes in white matter parameters which were located in the bilateral Inferior Fronto-Occipital Fascicle (IFOF), the Superior Longitudinal Fascicle (SLF), the Arcuate Fascicle (AF), the Uncinate Fascicle (UF) and the Corpus Callosum (CC) (13, 20, 22, 23, 33) which might be related to alterations in myelination or axonal characteristics (2, 34). White matter plasticity has been reported in relation to different aspects (i.e., novel speech sounds, vocabulary, grammar, etc.) of L2 acquisition (20, 24, 35) and respective variations across the phases of language learning (9, 10, 36).

Although it is widely accepted that language processing is dominated by the left hemisphere (31, 37), increasing evidence suggests that the right hemisphere is highly involved in L2 learning (10, 38, 39), including dynamic changes in lateralization across phases of language learning (40). In addition, differences in the CC have been reported between bilingual and monolingual participants, suggesting its involvement in L2 learning (9, 14). The CC is the structural bridge that allows the interaction between the hemispheres (41, 42). However, its role in the acquisition and use of a second language remains unclear.

There are two competing theories for the general role of the CC in interhemispheric interaction. One argues for the inhibition of the activation in the other hemisphere and the other suggests excitatory mechanisms (for reviews see (43, 44)). The interhemispheric inhibition theory proposes that an area can reduce the activity in homologous contralateral areas via the CC to allow for fast and automatic processing within each hemisphere and leading to functional hemispheric specialization. In contrast, the excitatory theory suggests that activation in one hemisphere facilitates the activation of homolog areas, increasing information exchange between the hemispheres. It is still an open question whether, during initial L2 learning, the role of the CC is mainly excitatory or inhibitory. However, it is well established that first language (L1) processing is strongly left lateralized and the CC establishes a strong inhibition from the dominant left hemisphere on the right hemisphere. In contrast, early phases of L2 learning involve the right hemisphere possibly due to weakened inhibition in early phases of L2 learning, hence allowing the engagement of the right homologs of the language areas. Accordingly, a decrease in CC-connectivity would follow a reduction of inhibitory explanation suggesting that this mechanism might play a role in L2 learning.

Here, we provide empirical data of longitudinal white matter changes as a function of L2 learning in two successive phases. Based on the DRM-model, we hypothesize that L2 learning-induced changes in structural connectivity will occur mainly after an initial beginner phase of learning, during which white matter changes are expected to be limited. In a second, intermediate consolidation phase, we expect changes in the structural connectivity of the language network. This learning phase involves both semantic processing of new vocabulary, and on the other hand, local syntactic processing based on lexical word category and semantic information. We hypothesize that these plastic changes take place primarily in the lexical-semantic system of both hemispheres and that such changes are related to the improvement in L2 performance. Furthermore, we hypothesize that L2 learning will lead to a significant change in transcallosal connectivity, supporting the role of the CC in interhemispheric communication.

To test these hypotheses, we recruited a large group of young, healthy Arabic native-speaking participants for an intensive German language course over six months to reach an intermediate proficiency (B1) level. The course consisted of an initial beginner phase of three months and a consolidation phase of the same duration. After three and six months, the participants took a standardized German language test that assessed L2 comprehension and production. At the beginning of the course and after each learning period, we acquired longitudinal high-resolution diffusion MR images and computed the white matter structural connectivity network in each participant. This structural network included the intrahemispheric connections between the language processing areas in the left hemisphere, and between their right hemisphere homologs as well as the callosal connections of these cortical areas in both hemispheres. Then, we compared the network properties between the different time points to identify changes in specific pathways and subnetworks. To test all connections for longitudinal changes in connectivity in an unbiased manner without introducing strong *a priori* hypotheses into the analysis, we used the recently proposed network-based statistics (NBS) (45) and a mixed-effects model (46). Changes in structural connectivity were then related to improvements in language tests to demonstrate a direct functional relevance of the detected changes in the language network.

## Results

### Improvement of L2 performance

The L2 performance after three months (59 participants) and after six months of learning (51 participants) was measured with standardized tests for German as an L2. The results of both tests were normalized to a common scale following the scaling method proposed in the Cambridge English Scale. Linear mixed-effects (LME) models were used in MATLAB to analyze behavioral improvement during learning, with time points modeled as a fixed effect (see Methods). The data showed a significant improvement in L2 performance between the two time points of the German tests (t=17.92, p<0.0001, see Figure 1A). After six months of learning, 41 participants took an additional vocabulary test. Correlation analysis between L2 vocabulary and the B1 language test showed that individuals with richer L2 vocabulary had higher overall language proficiency (r=0.509, p=0.002, see Figure 1B).

**Figure 1.**
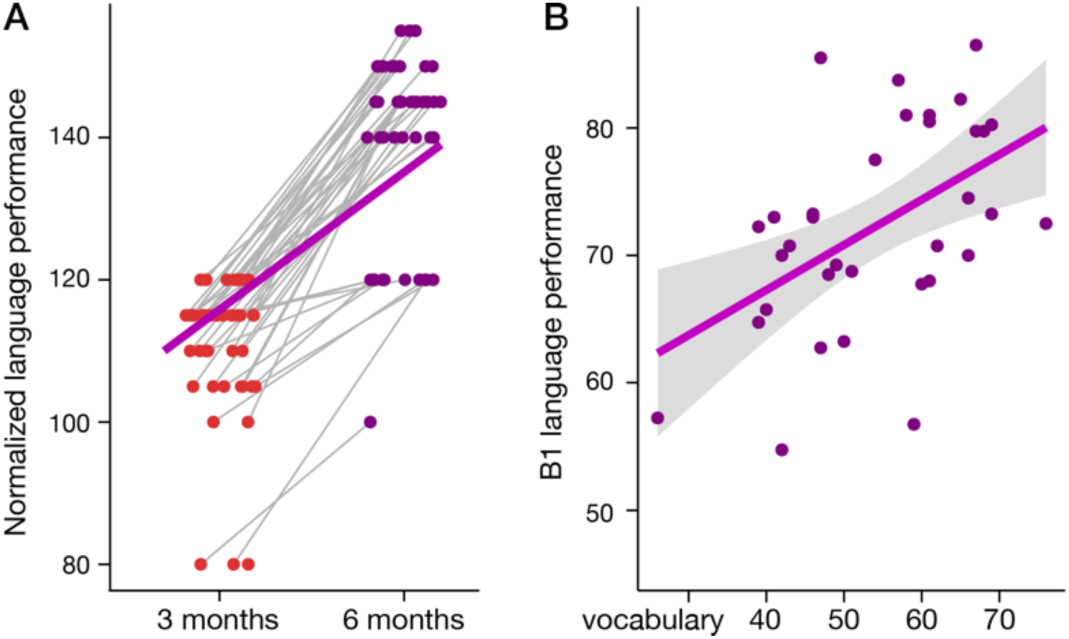
L2 improvement during learning. (A) Longitudinal changes of the normalized language performance after three and six months of L2 learning. (B) Correlation of L2 vocabulary score and overall language performance (B1 test) after six months of L2 learning.

### Lateralization and longitudinal changes of the intra- and interhemispheric connectivity

The initial lateralization test showed that the global intra-hemispheric connectivity in the language network is stronger in the left hemisphere than in the right hemisphere for each of the three measurement time points (baseline: left > right, t = 8.17, p < 0.0001; 3 months: left > right, t = 6.71, p < 0.0001; 6 months: left > right, t = 6.56, p < 0.0001; see Figure 2B). The longitudinal statistical analysis was then performed separately for the total intra-hemispheric connectivity in each side and the interhemispheric connectivity using an LME model with time points as a fixed effect. The result showed a significant dynamic decrease in interhemispheric connectivity during learning, specifically with an effect in the second half of the learning period (baseline-3 months: t = −0.56, p = 0.57 (n.s.); 3 months – 6 months: t = −7.33, p < 0.0001, baseline – 6 months: t = −8.97, p < 0.0001; see Figure 2C). However, we did not observe any significant changes in the longitudinal analysis of intra-hemispheric connectivity within the language network in each hemisphere or the lateralization index of the connectivity within the language network.

**Figure 2.**
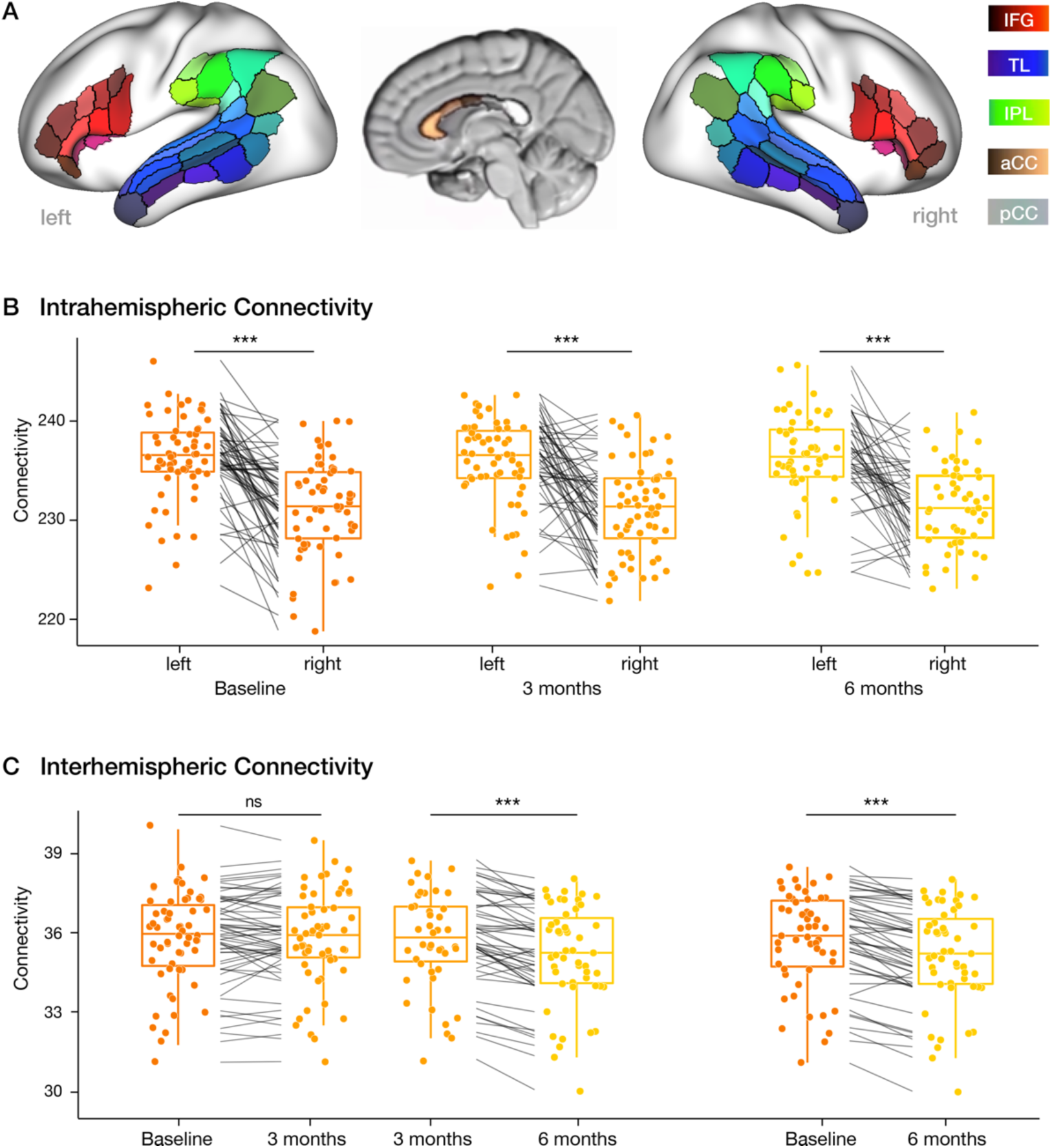
Longitudinal changes of the intra- and interhemispheric connectivity in the language network. (A) Areas in the language network of the left and right hemisphere in the inferior frontal gyrus (IFG), the inferior parietal lobe (IPL) the superior and middle temporal lobe (TL), and the anterior and posterior corpus callosum (aCC, pCC). (B) Intra-hemispheric connectivity at each time point during L2 learning shows significant left lateralization of the language network. (C) Longitudinal changes in interhemispheric connectivity show a significant decrease in the second learning period (middle) and over the full 6 months (right). The boxplots show the median, quartiles, 1.5* interquartile range, and all individual data points. (*** p<0.0001).

### Plasticity of the structural language subnetworks across different learning periods

The Network-Based R-Statistics (NBS) LME models (p-threshold = 0.01, K = 5000 permutations) revealed a complex reorganization of multiple subnetworks during L2 learning, including connections between all subregions in the bilateral temporal lobe (TL), inferior parietal lobe (IPL), and right inferior frontal gyrus (IFG, p < 0.01, NBS corrected, Supplementary Figure S2). To shed light on the temporal properties of network changes, post hoc LME analyses between adjacent time points allowed us to identify specific effects of each connectivity for the early and later learning period (p<0.05). During the first three months of learning, there was a significant decrease in connectivity for only a few connections belonging to three subnetworks. These subnetworks consisted of the interhemispheric connections of subregions of the IFG and parts of the right arcuate fascicle (AF) connecting the posterior IFG and the posterior middle and inferior temporal gyrus (Figure 3A). However, in the second learning period (from three to six months), the statistical analysis revealed an increase in connectivity in three subnetworks, including the bilateral parietal-temporal system as well as the right AF (see Figure 3B, left). Interestingly, the frontal and temporal-parietal interhemispheric networks showed decreased connectivity in this second period (see Figure 3B, right). The figure shows the mean changes of all individual connections within each subnetwork and the distribution of the changes. Individual data for each participant and each connection are shown in Supplementary Figure S4.

**Figure 3.**
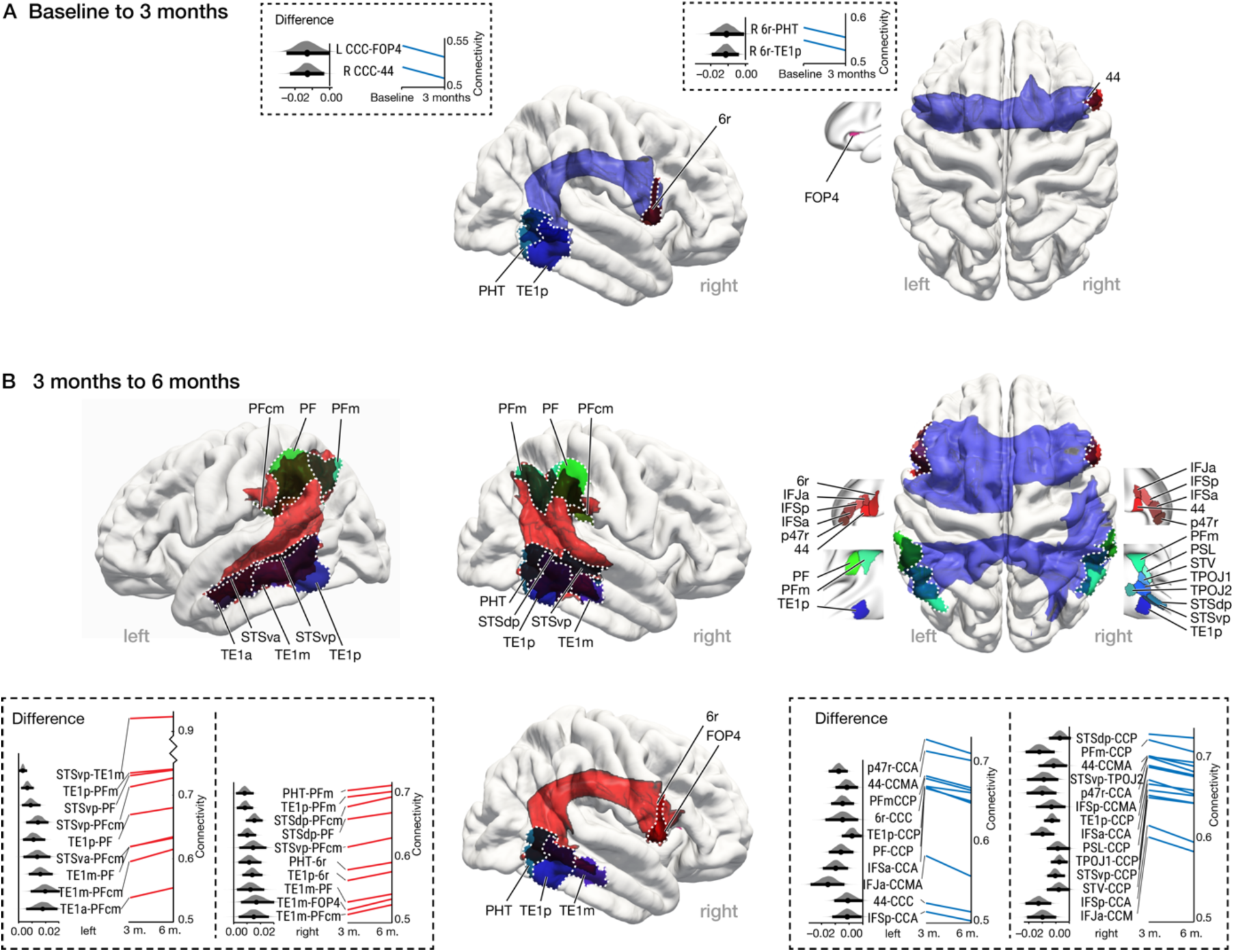
Subnetworks with longitudinally increased and decreased connectivity in the two learning periods. (A) First learning period: Decreasing connectivity (blue) in three small subnetworks including anterior transcallosal connections as well as the right AF. (B) Second learning period: Increasing connectivity (red) in three large subnetworks connecting the posterior temporal and the inferior parietal regions in both hemispheres along with the right AF (left). Decreasing connectivity (blue) in the anterior and posterior transcallosal subnetworks (right, all p<0.05 NBS corrected). The brain figure shows the group averaged probabilistic tractography of the subnetworks with increased (red) and decreased (blue) connectivity together with the corresponding brain regions. The figure in the box shows the effect size and change trend of each connection.

### Relationship between L2 proficiency and connectivity changes in the language network

To test the relationship between brain network plasticity and L2 performance increase over the different learning periods, we also used NBS with LME models. The initial behavioral analysis revealed that all participants showed an improvement in their L2 scores between three and six months of learning (Figure 1). This monotonic increase in performance allowed us to use a more parsimonious LME model that included only the L2 score and did not require modeling time as an additional separate factor. This NBS LME model allowed us to test for longitudinal correlations between the structural network characteristics and L2 performance for each participant after three and six months of learning. Figure 4 shows the brain subnetworks that show a significant linear relationship between the L2 score and the brain connectivity at three and six months (p<0.01, NBS corrected). The NBS LME analysis showed that the improvement in L2 proficiency was correlated with increased connectivity in subnetworks connecting the posterior temporal and the inferior parietal lobes in both hemispheres as well as in the right arcuate fascicle (AF) (see Figure 4A). Additionally, a negative correlation between connectivity changes and the L2 score was found in the anterior and posterior interhemispheric connections (p<0.01, NBS corrected, Figure 4B). In the frontal lobe, only the left subnetwork of the transcallosal connections showed a significant correlation. The figure shows the regression lines of the correlation for all individual connections within each subnetwork. The individual data for each participant and each connection are shown in Supplementary Figure S5.

**Figure 4.**
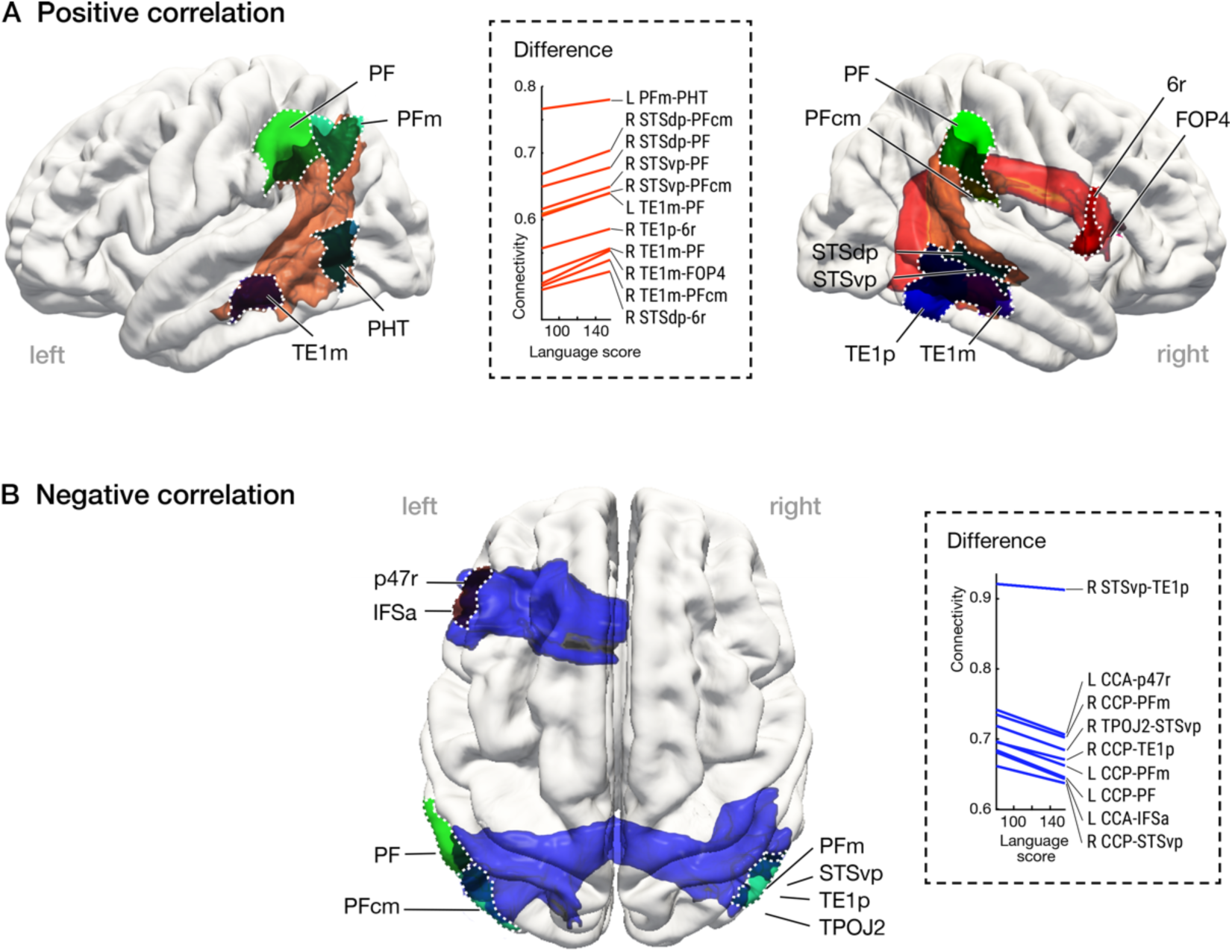
Relating changes in connectivity and the L2 proficiency from three to six months of L2 learning. (A) Positive correlation between L2 performance and connectivity changes in the left and right temporal-parietal network and the AF (red). (B) Negative correlation in the anterior (left) and posterior transcallosal network (blue, all p<0.05 NBS corrected). The plots display the regression lines for all connections in all networks. In the brain images, the colored mean tractography shows the correlated subnetworks together with the corresponding brain regions. The figure in the box shows the correlation trend between each connectivity and L2 proficiency.

Another measure of L2 learning success relates to L2 vocabulary size. In a *post hoc* analysis, we examined whether participants with a large L2 vocabulary showed different changes in white matter connectivity compared to those with a smaller L2 vocabulary. The group was divided based based on their productive vocabulary in the written text of the B1 test. After splitting into two groups, only the group with higher vocabulary scores showed a positive correlation between increased L2 scores and connectivity changes in the right temporal-parietal and AF subnetwork and a negative correlation with interhemispheric connectivity (p < 0.01, NBS corrected, Supplementary Figure S3). This suggests an important role of the right hemisphere for successful L2 learning.

## Discussion

The present longitudinal study tested the hypothesis that second language (L2) learning induces a dynamic reorganization of the structural white matter language network. For this purpose, Arabic native speakers participated in a six-month intensive language learning program in which they learned German as their L2. In the first half of the learning period (0 to 3 months), only small subnetworks with a few white matter connections showed significant changes in connectivity, whereas, in the second half of the learning period (3 to 6 months), significant changes were observed in multiple and larger white matter subnetworks. Specifically, the bilateral temporal-parietal system and the right arcuate fascicle (AF) showed increased connectivity. Additionally, the interhemispheric connectivity across the corpus callosum (CC) was reduced during this learning period. Most importantly, the brain changes in the late learning period correlated with the increases in L2 proficiency.

Our findings provide empirical evidence for the time course and location of white matter changes in the consolidation phase as suggested by the Dynamic Restructuring Model (DRM) of second language acquisition (15). This model suggests that plasticity in the white matter language network will emerge in a second phase of L2 acquisition, allowing for more efficient interaction between the different language areas within each hemisphere. This is indeed what we found in the present study.

### Dynamic intra-hemispheric white matter changes underlying L2 performance

The white matter network is the structural basis for neuronal communication between brain areas and its plasticity is crucial for learning new skills (2). Distinct subnetworks within the language system are specialized for different domains, and the corresponding connections are modulated by their usage. Therefore, we expected changes in the subnetworks reflecting specific tasks to be mastered in the newly learned language.

#### Semantic and phonological processes

The analysis of the language test showed a significant improvement in L2 from three to six months. L2 performance at six months correlated with the results of an independent L2 vocabulary test (47). From a neuroplasticity perspective, it is important to note that we found a significant change in the structural connectivity over this period in bilateral temporal-parietal subnetworks and right temporal-frontal connections. Crucially, these longitudinal changes in connectivity were found in subnetworks that were very similar to those that showed plastic changes correlated with L2 proficiency in our study. These subnetworks form the structural basis for lexical-semantic and phonological processing (31, 48, 49) and have previously been related to L2 vocabulary learning (24, 25, 50). In this regard, it is interesting to look at subgroups of participants with higher and lower L2 vocabulary scores. Our additional correlational analysis in these subgroups revealed a significant positive correlation between L2 improvement and changes in connectivity in the right fronto-temporal subnetwork as part of the AF in the second learning phase only in the group with a larger L2 lexicon compared to the other group (see Supplementary Figure S3). This result suggests that during this phase of L2 learning, the changes in the language network are related to the consolidation of lexical processing and highlights the importance of the right hemisphere for L2 acquisition.

Successful language learning depends on phonological discrimination during perception and phonological selection during production to decode speech sounds and associate them with the meaning of new words. In the neural language network, the bilateral inferior parietal lobe (IPL) and the superior temporal gyrus (STG) are involved in phonological storage and word decoding, and the middle temporal gyrus (MTG) is an integral region engaging in lexical-semantic access (31, 49, 51) Functional imaging studies of L2 learning (30) suggest a stronger functional activity and connectivity of the right IPL and STG during early learning phases, and phonological processing of L2 words (52, 53). Structural imaging studies demonstrate that these regions show changes in gray matter morphology during L2 learning that are related to L2 vocabulary acquisition and L2 proficiency (25, 26, 50, 52–55).

In addition to white matter effects in temporal-parietal connections, we also found increased connectivity in the right hemispheric temporal-frontal subnetwork as part of the AF which corresponds to the right hemispheric equivalent of the dorsal language network. This finding is consistent with previous studies highlighting the importance of the right hemisphere for L2 lexical-semantic and phonological processing during the initial and intermediate phases of adult L2 acquisition (10, 24). In addition, a higher involvement of the right prefrontal cortex (56) during L2 processing may be related to more working memory and attentional processes in L2 (57). Taken together, our findings strongly suggest that efficient L2 learning, especially vocabulary acquisition in adults, involves the right fronto-parieto-temporal network.

### Longitudinal decrease in transcallosal interhemispheric connectivity

The observed network strength in both hemispheres showed that the language network is lateralized to the left at all measured time points during L2 learning. This is in line with the widely accepted model that the language network is dominated by the left hemisphere (31). In previous studies (10, 32), and the present data, there is evidence of increased right hemisphere involvement during L2 learning, as reflected by strong changes in white matter connectivity in the right hemisphere. These changes might be directly related to the reduced transcallosal connectivity, allowing for additional L2 processing to occur in the right hemisphere (43). Indeed, we found a significant decrease in the interhemispheric connectivity in the anterior and posterior CC during L2 learning. This reduction correlated with the increase in L2 performance in the second learning phase. The current data provide a comprehensive demonstration of the role of the CC in L2 learning. In native language processing, the dominant left hemisphere exerts an inhibitory influence on the non-dominant right hemisphere via the CC (44). However, during the initial and intermediate phases of L2 learning, a highly involved right hemisphere language network is required to build up the L2 lexicon. This would explain why successful L2 acquisition is accompanied by a decrease in transcallosal connectivity. This reduces the inhibition of the language-dominant left hemisphere on the corresponding regions in the right hemisphere, allowing increased processing and connectivity to occur in the right half of the brain. However, this fundamentally new finding needs to be further explored and supported by additional data.

## Conclusions

Our study showed that L2 learning in adults leads to dynamic changes in brain connectivity within and across hemispheres. The experimental evidence suggests that plastic changes in the white matter system occur mainly after an initial period of learning. In this phase, the adaptation of the language network appears to be focused on the lexical-semantic system, particularly in the temporal and temporal-parietal regions, with increased connectivity in each of the two hemispheres and strong involvement of the right side. At the same time, L2 learning leads to reduced connectivity between the hemispheres, which could result in reduced inhibition of the dominant left hemisphere on the right, temporarily freeing up resources in the right half of the brain for efficient learning of new L2 linguistic features.

## Materials and Methods

### Participants

Eighty-four young, healthy right-handed Arabic native speakers were recruited for an intensive German course (5 h/day, 5 days/week) over six months to reach the threshold level B1 according to the Common European Framework of Reference for Languages (CEFR, (58)). The course was divided into two learning phases of three months each. During the two learning periods, some participants left the language course for personal reasons and were not included in the corresponding analysis. Fifty-nine participants completed the first learning phase (mean age, 24.4 ± 4.5 (SD) years, 51 male) and 51 participants completed the two learning phases (mean age, 24.7 ± 4.6 (SD) years, 43 male). After each learning phase, participants took a 90-minute standardized second language proficiency test (A1 and B1 tests of the Goethe Institute). After six months of learning, an additional L2 Vocabulary Size Test (VST) was taken by a subgroup of 41 participants (35 male). All participants were immersed in the second language environment and lived in Germany during the course. All participants were native speakers of the Levantine dialect of Arabic and of normal intelligence (non-verbal Raven’s matrix test (59), score 50.4 ± 6.7, ranging around the upper 90 percentile of the reference population, subgroup N=32) and spoke only one native language. All participants were recruited in Leipzig and arrived in Germany 6-8 months before the start of the study. They were settled in Leipzig for a long-term stay and were highly motivated to learn German and integrate into the academic system. An initial German test revealed that the entire group had no-to-minimal knowledge of German, well below beginner level A1. Structural and high-angular and spatial resolution diffusion MRI data were acquired from each participant on a Siemens 3T Prisma MRI scanner at baseline and after three and six months of learning. Details of the learning procedure and MRI acquisition can be found in the Supplementary Materials. The study was approved by the Ethics Committee of the University of Leipzig, and all participants gave written informed consent in their native language.

### Structural language connectome

We used probabilistic diffusion MRI tractography to compute the white matter network between the language processing regions (Figure 2A) in each participant and time point (baseline, three and six months of learning). The analysis followed the previously established method (7). Cortical seed and target areas were defined using the Human Connectome Project (HCP) fine-grained atlas in addition to a subdivision of the corpus callosum (CC) atlas (60, 61). The core regions of the language network, as defined previously (31), included the dorsal and ventral pathways in both hemispheres between subregions in the bilateral inferior frontal gyrus (IFG), superior temporal gyrus (STG), middle temporal gyrus (MTG), and inferior parietal lobe (IPL). To account for interhemispheric connections, we included white matter regions in the medial cross-section of the CC resulting in 33 cortical and 5 CC regions per hemisphere (see Figure 2A and Supplementary Materials, Table S2). The CC is a bottleneck for estimating interhemispheric connections, and direct one-to-one connectivity between cortical areas in both hemispheres cannot be robustly estimated by tractography. Therefore, we computed probabilistic tractography between cortical and CC regions as a robust approximation of the interhemispheric connectivity. To remove false-positive connections (62), we retained the 30% strongest connections for the network analysis. Details of the connectivity analysis can be found in the Supplementary Materials.

### Statistical analyses

To estimate longitudinal changes in the behavioral learning progress, a linear mixed-effects model (LME) (i.e., y ~ time + (1 | participant); y represents L2 proficiency) with time points as the fixed effect and participant as the random effect was applied to the scaled language test scores obtained after three and six months of learning to analyze changes across learning periods. In addition, a correlation analysis was performed between the L2 vocabulary level after six months and the B1 language test scores to analyze how strongly the composed language test is related to vocabulary knowledge at this stage in this group.

To assess the relationship between L2 learning and plasticity in the white matter language network, we first tested the changes in the overall network strength within and between brain hemispheres. We first measured the network strength in each hemisphere (sum of all connectivity values between the cortical regions) and compared this parameter between hemispheres at each time point (baseline, three and six months of learning) to test for lateralization of the language network using a paired t-test. Next, we used a separate LME models (i.e., y ~ time + (1 | participant); y represents connectivity) with the three measurement time points as a fixed effect to test longitudinal learning-induced changes in interhemispheric connectivity as well as intra-hemispheric changes in the language network within each hemisphere (left and right) and the lateralization index. The interhemispheric network strength is the sum of the weighted connections between all cortical language regions and the CC, representing the connections crossing to the other hemisphere. In all analysis steps discussed so far, the LME models used time as a fixed effect and participant as a random effect, and the statistical tests were performed in MATLAB.

To localize subnetworks showing longitudinal changes within the language network across all time points, the recently proposed network-based R-statistics (NBS) (45) for LME models (46) with time points as a fixed effect and participant as a random effect was used. In the first step, the connectivity measures at different time points were modeled as fixed effects and participants were entered as random effects. This allowed us to analyze the longitudinal change of the structural connectome across the three different time points and account for individual differences (i.e., y ~ time + (1| participant); y represents connectivity). Then, posthoc statistics were used to identify subnetworks with significant changes between each pair of measurement points and to determine in which of the two language learning phases white matter changes occurred. Finally, to test whether changes in connectivity were related to individual L2 performance and to localize such subnetworks, we employed a second type of model and included L2 proficiency test scores as a fixed effect and participant as a random effect in the NBS LME models, allowing us to account for the longitudinal interactions between brain structural connectivity and L2 proficiency (i.e., y ~ score + (1|participant); y represents connectivity). Since test scores could only be acquired after three and six months of learning, these two time points were considered in this analysis. To visualize the statistically identified subnetworks, additionally, probabilistic tractography was computed between the regions belonging to these subnetworks. The individual pathway maps were normalized, averaged, and visualized together with the regions of the subnetwork. This allowed to show the white matter pathways belonging to the identified subnetworks.

## Acknowledgments

We would like to thank S. Brogatio of the former Leipzig Refugee Council Leipzig for her help in recruiting the participants as well as all participants, teachers, and student helpers for their contributions. We thank Prof. C. Fandrych, Herder Institute, Univ. Leipzig for his support in the conception and design of the language courses, and Prof. E. Tschirner, ITT, Univ. Leipzig, for his support in the language tests, especially in the application of the vocabulary tests. Finally, we would like to thank M. Lišaník for his contribution to data acquisition, participant testing, and study conception. This work was supported by the SPP2041 program “Computational Connectomics” of the German Research Foundation (DFG), grant number 347141397 (AN 1156/1-1, FR 519/22-1).

## Supplementary Materials

### Supplementary Methods

#### Language learning procedure

In our study, we recruited a large group of young, healthy, native Arabic speakers to participate in a six-month intensive German language course to reach an intermediate level of proficiency (B1, first level of independent language proficiency). The second language (L2) teaching and assessment structure followed the Common European Framework of Reference for Languages (1, 2). According to the CEFR, the first intermediate level of independent language use (B1) can be reached after approximately 600 hours of language learning, which corresponds to a six-month intensive course. The courses have been structured in cooperation with the Herder-Institute of the University of Leipzig, Germany, which is specialized in research and teaching of German as an L2. The proficiency levels of this standard framework comprise six levels. Levels A1 and A2 represent the elementary use of the language for beginners. Levels B1 and B2 represent intermediate language levels. At the first intermediate level (B1) of German, a learner can understand the main points when clear, standard language is used and familiar topics related to work, school, leisure time, etc. are the focus. The learner can make a short statement to explain his views and plans. C1 and C2 are the highest possible levels. In our study, the participants underwent two phases (0-3 months and 3-6 months) of daily intensive classroom training in German (L2). The course combined classroom teaching using standard textbooks, complex naturalistic speaking and reading, and clear instruction in grammar and vocabulary. The course took place at the Max Planck Institute in Leipzig, in small groups of 12-15 students, 45 minutes per lesson, 5 lessons per day, 5 days per week. Three different professional teachers taught the classes in each group to increase the language input of the learners and to reduce instructional variance between the groups. Daily homework was assigned by the teachers and consisted mainly of consolidating and reviewing the topics taught. In addition, several Arabic-speaking student assistants helped the learners with everyday issues so that they could concentrate fully on the language courses.

#### Language proficiency test

After 3 months and 6 months of learning, participants took a 90-minute standardized second language proficiency test that assessed language comprehension and production performance through four subtests in listening, reading, writing, and speaking in German. Language acquisition was assessed in the first phase between 0 and 3 months with the standardized A1 test and in the second phase between 3 and 6 months with the B1 language test of the German Goethe Institute, which tests oral and written, receptive and productive skills (i.e. listening, reading, speaking, and writing). An additional L2 Vocabulary Size Test (VST) was taken by a subgroup of 41 participants (35 males) after 6 months of learning (3). This receptive VST was developed at the Institute for Test Research and Test Development, Leipzig (ITT-Leipzig, http://www.itt-leipzig.de, (4)). The students were tested on their knowledge of a sample of the 3000 most common words required at this level. It measures in five sections, how many of a sample of words belonging to a given frequency range are known (1000 most frequent, 2000 most frequent, etc.). This results in a maximum score of 30 points per section, which were added together.

### Transformation of the A1 and B1 language scores to a common scale

To estimate the L2 proficiency longitudinally and correlate it with the brain structural plasticity during learning, scores from each language test at each time point were scaled to a common scale following the Cambridge English Scale (https://www.cambridgeenglish.org/exams-and-tests/cambridge-english-scale). In our study, the progress scale was always divided into steps of 5. The detailed conversion relationship between the test score and the common scale is shown in Supplementary Table S1.

#### MRI data acquisition

Structural and high-resolution diffusion-weighted MR images were acquired on a 3 Tesla Prisma MRI system (Siemens Healthineers, Erlangen, Germany) with a 32-channel head coil with the following scanning parameters: Isotropic voxels resolution of 1.3 mm, 60 diffusion directions (b = 1000 s/mm^2^) and 7 images without diffusion weighting (b = 0 s/mm^2^), TE = 75 ms, TR = 6 s, GRAPPA = 2, CMRR-SMS=2, 3 repetitions to improve the signal-to-noise ratio, and 2 b0 acquisitions with opposite phase encoding. The diffusion sequence was repeated 3 times to increase the. For the anatomical segmentation, we acquired quantitative multiparametric structural images with 1 mm resolution (5). The images were preprocessed using the publicly available hMRI toolbox (http://hmri.info) and the quantitative magnetization transfer (MT) images were used for the segmentation and parcellation steps.

### Connectivity analysis

#### Diffusion MRI preprocessing

Preprocessing of diffusion data was performed using the FMRIB Software Library (FSL, http://www.fmrib.ox.ac.uk/fsl). Diffusion images were corrected for susceptibility and eddy current induced distortions as well as head motion with the FSL tools “topup” and “eddy” using optimized parameters matched to image resolution. Imaging noise in the high-quality diffusion MRI data was minimized by combining the three repetitions. No additional denoising algorithms were applied to minimize image blurring. The optimized imaging settings allowed Gibbs ringing artifacts to be minimized and an additional correction was not required. No additional intensity or bias field correction was applied. Finally, the brain volume was masked from the background and the standard DTI contrast maps were computed. The processed datasets were checked individually to exclude artifacts from the acquisition or preprocessing. Finally, the voxel-wise fiber distribution for probabilistic tractography was computed with up to 3 fiber directions per voxel using the FSL command “bedpostX” (6).

#### Surface segmentation

For each participant, the structural connectome of each hemisphere was computed as follows (see also Supplementary Figure S1). First, the cortical and white matter surface, as well as the 5 subsections of the corpus callosum of each participant were generated from the MT images using FreeSurfer 5.3 (http://surfer.nmr.mgh.harvard.edu, (7). The white matter surface was shifted 1 mm into the white matter using the FreeSurfer command “mris_expand” to define robust seed and target regions for probabilistic tractography.

#### Parcellation of the seed regions

The cortical surface was divided into 180 regions in each hemisphere using the multi-modal parcellation developed as part of the Human Connectome Project (HCP) (8). Therefore, the atlas annotations were transformed into separate labels using “mri_annotation2label“, and then mapped to each participant using “mri_label2label”. Finally, the labeled white matter surface (shifted 1mm inside the white matter) was mapped to the individual anatomical voxel space using “mri_label2vol” to generate a voxel-based definition of parcellation corresponding to the cortical areas. The labeled cortical and CC regions were registered to the diffusion images (FA contrast) using a rigid body registration using “flirt” and applied to the labeled regions using nearest-neighbor interpolation. Using the corpus callosum sections as seed and target regions in probabilistic tractography reduces the problem of spurious white matter connections resulting from the tracking through the bottleneck of the corpus callosum. This results in a more robust estimation of the inter-hemispheric connectivity.

#### Connectivity estimation

All registered regions were used as seed areas for probabilistic tractography (6) using “probtrackX” with default parameters. The structural connectivity between all regions was computed representing the relative number of streamlines between all pairs of regions. Those connectivity estimates are influenced by the local microstructural properties of the pathway between the regions and integrate the local properties into one connectivity estimate for a specific connection. The estimated connectivity values for all regions (full HCP atlas) were logarithmically scaled and normalized by the size of the seed region (log of the number of seeded streamlines) to build a connectivity matrix with normalized values ranging from zero to one. Next, the values for both tracking directions were averaged (from region A to region B, and from region B to region A). For each participant and each hemisphere, we obtained the weighted symmetric connectome matrix (Supplementary Figure S1).

#### Network thresholding

Additionally, we removed weak and noisy connections below a predefined threshold (in the average matrix across all participants) as they cannot be estimated reliably with tractography due to the limited sampling of the distribution that may result in false-positive connections (9). This allowed the exclusion of connections that did not align with the major fiber pathways in the human brain (10), and removed, e.g., connections between the left parietal lobe and frontal CC regions that do not exist anatomically. To determine this threshold, we increased the threshold in increments of 10% to create seven networks with different densities. Network thresholding methods were shown to be able to disentangle spurious and genuine connections (9). These networks ranged from a dense network that contained 80% of all connections to a sparse network that included only the strongest 20% per hemisphere. A threshold of 30% was found to reliably remove implausible false-positive connections and still retain the major pathways for the network-based analysis. With this global density threshold, 67% of the connections within the cortical language network (out of 528 per hemisphere, 33*32/2) and 55% of all possible connections including the CC areas were retained. The same network mask was applied to every individual participant. Finally, the 33 language ROIs in each hemisphere were selected and the matrix was reduced to those elements for further analysis.

#### Network-based statistics

We used network-based statistics (NBS) (11) to identify subnetworks with systematic structural changes. NBS is a method to control for the family-wise error rate when testing each connection in the network by using the extent to which the edges are connected. Therefore, all connected components that were present in the set of supra-threshold connections (T-threshold = 3.3) were identified and the number of connections was stored. To estimate the significance of each component, NBS performed a nonparametric permutation test (K = 5000 permutations). At each permutation, the group to which each participant belonged was randomly exchanged, the same threshold is applied to create the set of connections above the threshold for each K permutation, and then the statistical test was recalculated and the size of the largest component m in the set of supra-threshold connections was stored. The p-value of each connected component of size m was then estimated by searching for the proportion of permutations for which the maximal component size was greater than m, and was then normalized by K. In this way, the NBS attempts to utilize the presence of any structure exhibited by the connections comprising the effect or contrast of interest to yield greater power than what is possible by independently correcting the p-values computed for each link using a generic procedure to control the FWE (11, 12).

### Supplementary Figures

**Figure S1:**
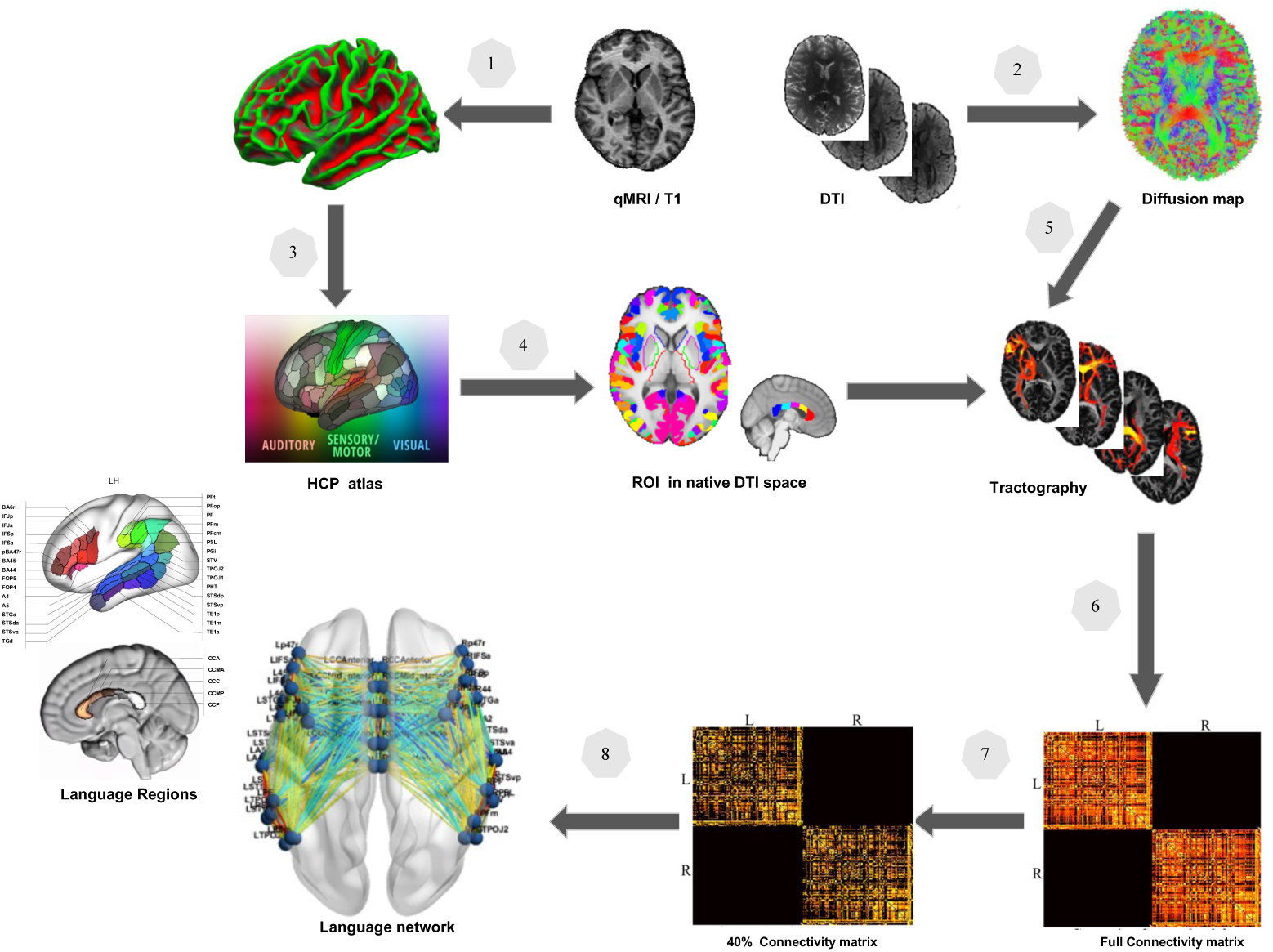
Processing pipeline of structural connectome construction. 1. White matter surface was generated by a segmentation of the anatomical MRI. 2. Preprocessing of the diffusion MRI. 3. Atlas-parcellation of cortical regions in the native brain surface. 4. Registration of the parcellated surface to native diffusion space. 5. Tractography using seed regions in diffusion space. 6. Whole brain network computation. 7. Network thresholding. 8. Extraction of the language network.

**Figure S2:**
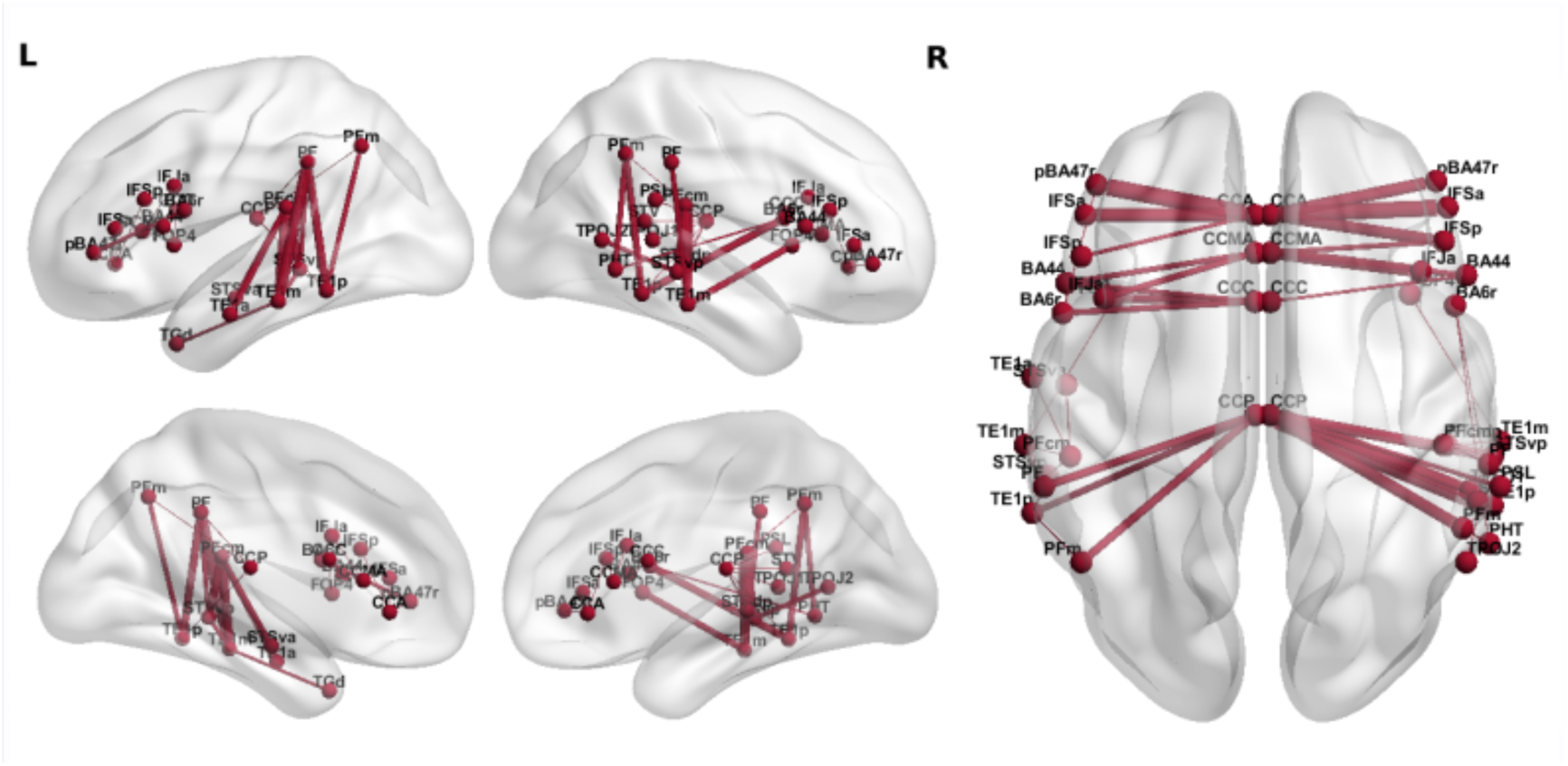
Longitudinal network changes across 3 learning time points. (p<0.05 NBR corrected).

**Figure S3:**
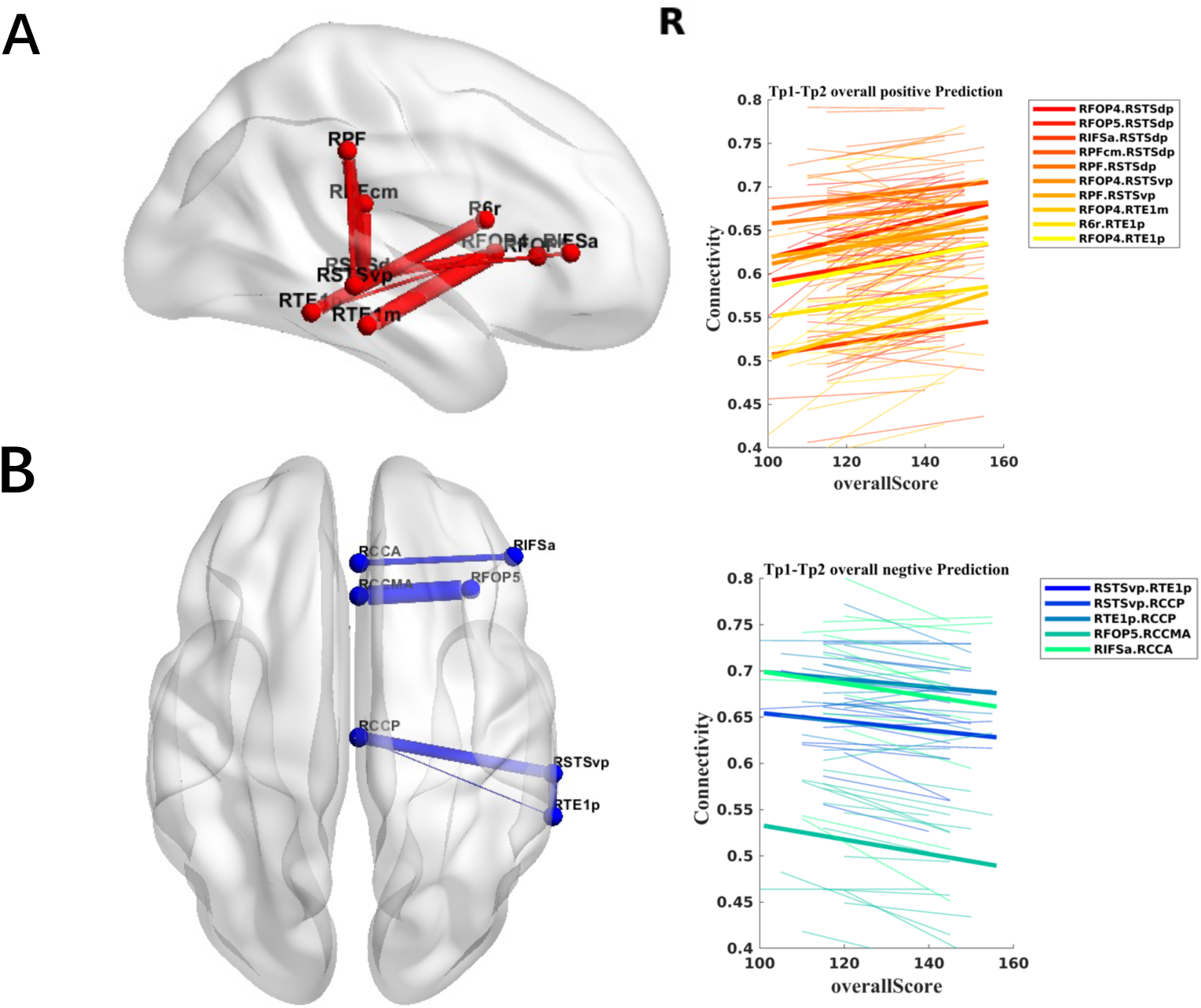
Subgroup with high productive vocabulary scores in the B1 test. The correlation between the changes in connectivity and L2 proficiency from three to six months of L2 learning. Positive (A) and negative (B) correlation between L2 performance and changes in subnetworks. Note that the connection between the right temporal lobe and the frontal lobe follows the right arcuate fascicle as shown in Figure 3 of the main manuscript.

**Figure S4:**
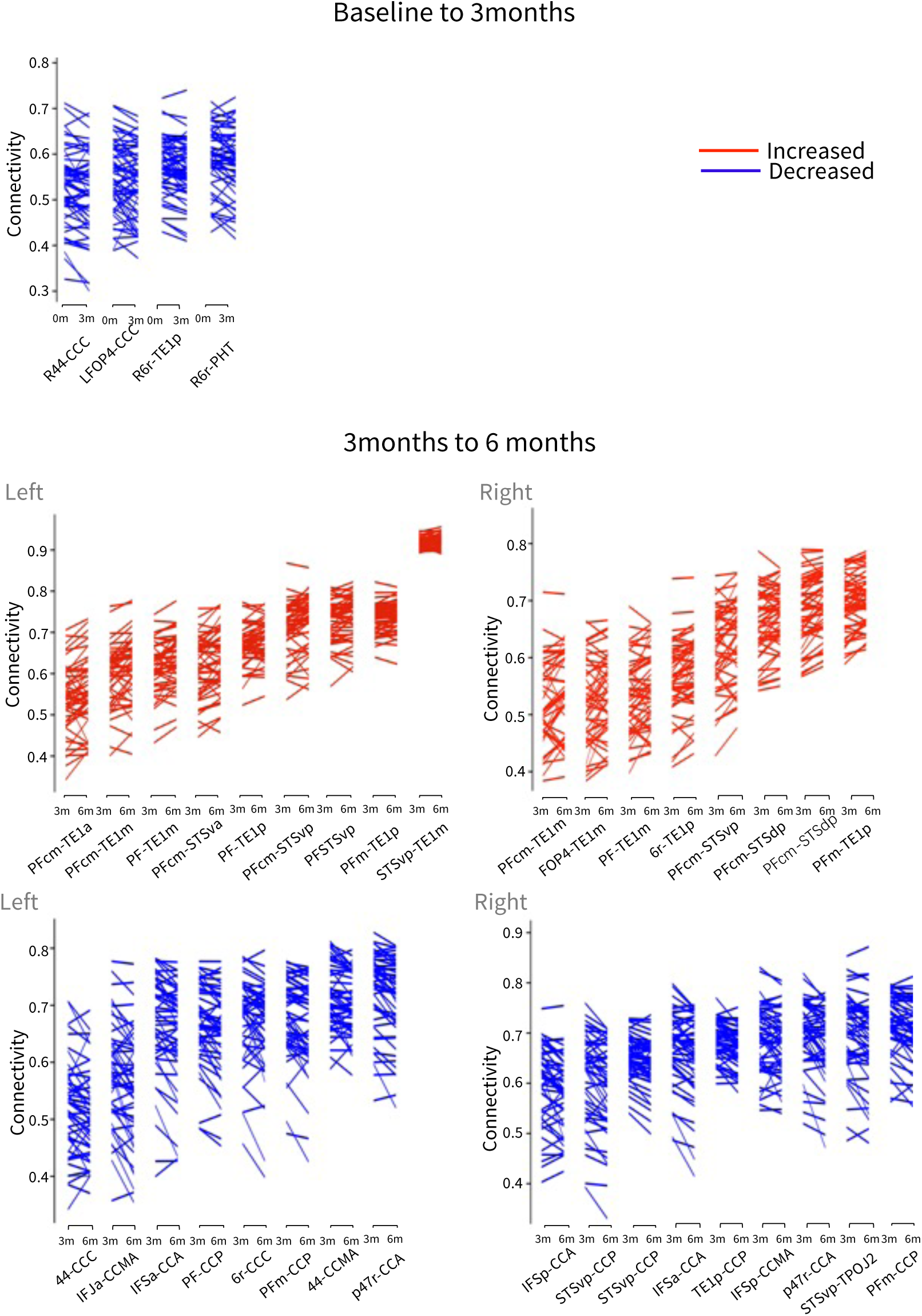
Individual data. Subnetworks with longitudinal increased and decreased connectivity in different L2 learning periods of each participant and each connection within the networks which showed significant changes.

**Figure S5:**
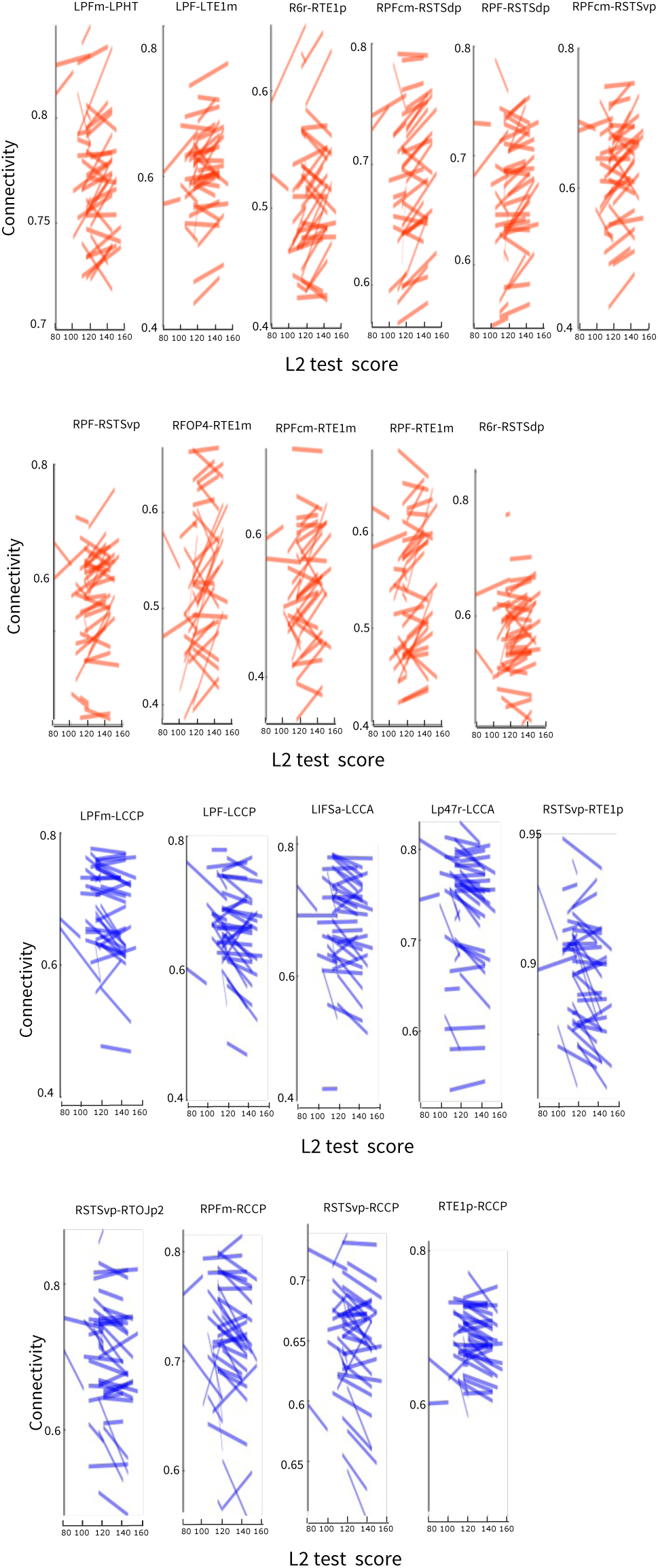
Individual data. Longitudinal changes of the connectivity values in relation to the progress in the language test from 3 months to 6 months of learning (normalized scores).

**Supplementary Table 1:**
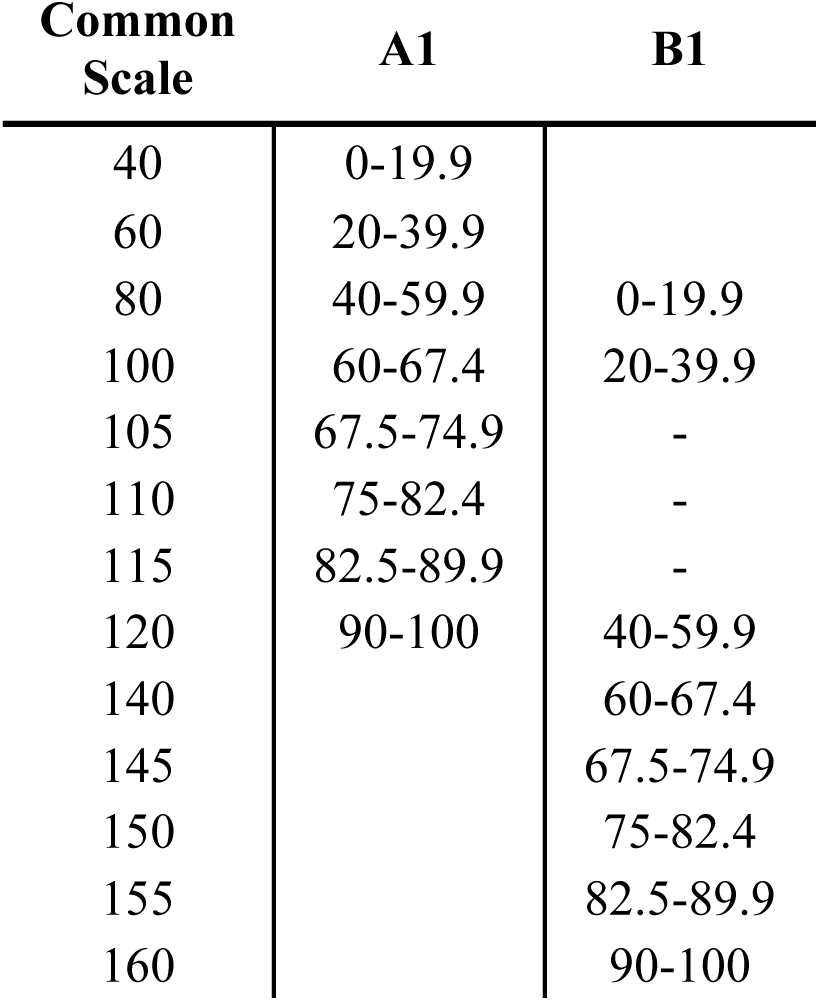
Transformation of the A1 and B1 language test scores to a common scale. The standardized tests require 60% to successfully pass the exam and are less quantitative below this threshold. Therefore, the conversion follows coarser discretization steps of 10 below 60%.

| Common<br>Scale | A1 | B1 |
| --- | --- | --- |
| 40 | 0-19.9 |  |
| 60 | 20-39.9 |  |
| 80 | 40-59.9 | 0-19.9 |
| 100 | 60-67.4 | 20-39.9 |
| 105 | 67.5-74.9 | - |
| 110 | 75-82.4 | - |
| 115 | 82.5-89.9 | - |
| 120 | 90-100 | 40-59.9 |
| 140 |  | 60-67.4 |
| 145 |  | 67.5-74.9 |
| 150 |  | 75-82.4 |
| 155 |  | 82.5-89.9 |
| 160 |  | 90-100 |

**Supplementary Table 2:** Labels for each language region. Inferior Frontal Gyrus: IFG, Temporal Lobe: TL, Inferior Parietal Lobe: IPL, anterior/posterior Corpus Callosum: aCC / pCC.

| HCP Atlas | ROI Name | Area Description | Region |
| --- | --- | --- | --- |
| 4 | BA44 | Broca Area 44 | IFG |
| 75 | BA45 | Broca Area 45 | IFG |
| 78 | BA6r | rostral Broca Area 6 | IFG |
| 79 | IFJa | anterior Inferior Frontal Junction | IFG |
| 80 | IFJp | posterior Inferior Frontal Junction | IFG |
| 81 | IFSp | posterior Inferior Frontal Sulcus | IFG |
| 82 | IFSa | anterior Inferior Frontal Sulcus | IFG |
| 108 | FOP4 | Frontal OPercular area 4 | IFG |
| 169 | FOP5 | Frontal OPercular area 5 | IFG |
| 171 | pBA47r | posterior Broca Area 47 rostral | IFG |
| 76 | 47l | Broca Area 47 lateral | IFG |
| 105 | PFcm | Area PFcm | IPL |
| 116 | PFt | Area PFt | IPL |
| 147 | PFop | Area PF opercular | IPL |
| 148 | PF | Area PF Complex | IPL |
| 149 | PFm | Area PFm Complex | IPL |
| 150 | PGi | Area PGi | IPL |
| 25 | PSL | PeriSylvian Language area | IPL |
| 140 | TPOJ2 | Temporo Parieto Occipital Junction 2 | IPL |
| 123 | STGa | Superior Temporal Gyrus anterior | TL |
| 125 | A5 | Auditory 5 Complex | TL |
| 128 | STSda | Superior Temporal Sulcus dorsal anterior | TL |
| 129 | STSdp | Superior Temporal Sulcus dorsal posterior | TL |
| 137 | PHT | Area PHT | TL |
| 175 | A4 | Auditory 4 Complex | TL |
| 176 | STSva | Superior Temporal Sulcus ventral anterior | TL |
| 130 | STSvp | Superior Temporal Sulcus ventral posterior | TL |
| 131 | TGd | Temporal pole dorsal | TL |
| 132 | TE1a | Temporal area 1 anterior | TL |
| 177 | TE1m | Temporal area 1 middle | TL |
| 133 | TE1p | Temporal area 1 posterior | TL |
| 139 | TPOJ1 | Temporo Parieto Occipital Junction 1 | TL |
| 28 | STV | Superior Temporal Visual area | TL |
| -- | CCa | Corpus Callosum anterior | aCC |
| -- | CCma | Corpus Callosum middle anterior | aCC |
| -- | CCc | Corpus Callosum central | aCC |
| -- | CCmp | Corpus Callosum middle posterior | pCC |
| -- | CCp | Corpus Callosum posterior | pCC |

## Notes

**Competing Interest Statement:** The authors declare no conflict of interest.

### Competing Interest Statement

The authors have declared no competing interest.

## References

1. R. D. Fields, White matter in learning, cognition and psychiatric disorders. Trends Neurosci. 31, 361–370 (2008).

2. R. J. Zatorre, R. D. Fields, H. Johansen-Berg, Plasticity in gray and white: neuroimaging changes in brain structure during learning. Nat. Neurosci. 15, 528–536 (2012).

3. B. Draganski, G. Christian, B. Volker, S. Gerhard, B. Ulrich, M. Arne, Changes in grey matter induced by training. Nature. 427, 311–312 (2004).

4. E. A. Maguire, D. G. Gadian, I. S. Johnsrude, C. D. Good, J. Ashburner, R. S. J. Frackowiak, C. D. Frith, Navigation-related structural change in the hippocampi of taxi drivers. Proc. Natl. Acad. Sci. U. S. A. 97, 4398–4403 (2000).

5. J. Scholz, M. C. Klein, T. E. J. Behrens, H. Johansen-berg, Training induces changes in white matter architecture. Nat. Neurosci. 12, 1370–1371 (2009).

6. M. Taubert, B. Draganski, A. Anwander, K. Muller, A. Horstmann, A. Villringer, P. Ragert, Dynamic properties of human brain structure: learning-related changes in cortical areas and associated fiber connections. J. Neurosci. 30, 11670–11677 (2010).

7. X. Wei, H. Adamson, M. Schwendemann, T. Goucha, A. D. Friederici, A. Anwander, Native language differences in the structural connectome of the human brain. Neuroimage. 270, 119955 (2023).

8. Z. Qi, M. Han, K. Garel, E. San Chen, J. D. E. Gabrieli, White-matter structure in the right hemisphere predicts mandarin chinese learning success. J. Neurolinguistics. 33, 14–28 (2015).

9. P. Li, J. Legault, K. A. Litcofsky, Neuroplasticity as a function of second language learning: anatomical changes in the human brain. Cortex. 58, 301–324 (2014).

10. Z. Qi, J. Legault, Neural hemispheric organization in successful adult language learning: Is left always right? Psychol. Learn. Motiv. 72, 119–163 (2020).

11. G. Videsott, B. Herrnberger, K. Hoenig, E. Schilly, J. Grothe, W. Wiater, M. Spitzer, M. Kiefer, Speaking in multiple languages: neural correlates of language proficiency in multilingual word production. Brain Lang. 113, 103–112 (2010).

12. A. Hahne, A. D. Friederici, Processing a second language: Late learners’ comprehension mechanisms as revealed by event-related brain potentials. Biling. Lang. Cogn. 4, 123–141 (2001).

13. A. A. Schlegel, J. J. Rudelson, P. U. Tse, White matter structure changes as adults learn a second language. J. Cogn. Neurosci. 24, 1664–1670 (2012).

14. P. E. Coggins, T. J. Kennedy, T. A. Armstrong, Bilingual corpus callosum variability. Brain Lang. 89, 69–75 (2004).

15. C. Pliatsikas, Understanding structural plasticity in the bilingual brain: The Dynamic Restructuring Model. Biling. Lang. Cogn. 23, 459–471 (2020).

16. H. Clahsen, C. Felser, How native-like is non-native language processing? Trends Cogn. Sci. 10, 564–570 (2006).

17. S. A. Rüschemeyer, C. J. Fiebach, V. Kempe, A. D. Friederici, Processing lexical semantic and syntactic information in first and second language: FMRI evidence from German and Russian. Hum. Brain. Mapp. 25, 266–286 (2005).

18. E. Higby, J. Kim, L. K. Obler, Multilingualism and the brain. Annu. Rev. Appl. Linguist. 33, 68–101 (2013).

19. I. Wartenburger, H. R. Heekeren, J. Abutalebi, S. F. Cappa, A. Villringer, D. Perani, Early setting of grammatical processing in the bilingual brain. Neuron. 37, 159–170 (2003).

20. P. K. Kuhl, J. Stevenson, N. M. Corrigan, J. J. F. van den Bosch, D. D. Can, T. Richards, Neuroimaging of the bilingual brain: Structural brain correlates of listening and speaking in a second language. Brain Lang. 162, 1–9 (2016).

21. S. Murray Sherman, The thalamus is more than just a relay. Curr. Opin. Neurobiol. 17, 1–7 (2007).

22. P. C. Mamiya, T. L. Richards, B. P. Coe, E. E. Eichler, P. K. Kuhl, Brain white matter structure and COMT gene are linked to second-language learning in adults. Proc. Natl. Acad. Sci. U. S. A. 113, 7249–7254 (2016).

23. C. Pliatsikas, E. Moschopoulou, J. D. Saddy, The effects of bilingualism on the white matter structure of the brain. Proc. Natl. Acad. Sci. U. S. A. 112, 1334–1337 (2015).

24. C. Hosoda, K. Tanaka, T. Nariai, M. Honda, T. Hanakawa, Dynamic neural network reorganization associated with second language vocabulary acquisition: a multimodal imaging study. J. Neurosci. 33, 13663–13672 (2013).

25. J. Legault, S. Y. Fang, Y. J. Lan, P. Li, Structural brain changes as a function of second language vocabulary training: Effects of learning context. Brain Cogn. 134, 90–102 (2019).

26. J. Mårtensson, J. Eriksson, N. C. Bodammer, M. Lindgren, M. Johansson, L. Nyberg, M. Lövdén, Growth of language-related brain areas after foreign language learning. Neuroimage. 63, 240–244 (2012).

27. F. M. Richardson, M. S. C. Thomas, R. Filippi, H. Harth, C. J. Price, Contrasting effects of vocabulary knowledge on temporal and parietal brain structure across lifespan. J. Cogn. Neurosci. 22, 943–954 (2010).

28. E. B. Barbeau, X. J. Chai, J. K. Chen, J. Soles, J. Berken, S. Baum, K. E. Watkins, D. Klein, The role of the left inferior parietal lobule in second language learning: An intensive language training fMRI study. Neuropsychologia. 98, 169–176 (2017).

29. S. Caffarra, N. Molinaro, D. Davidson, M. Carreiras, Second language syntactic processing revealed through event-related potentials: an empirical review. Neurosci. Biobehav. Rev. 51, 31–47 (2015).

30. K. M. Tagarelli, K. F. Shattuck, P. E. Turkeltaub, M. T. Ullman, Language learning in the adult brain: A neuroanatomical meta-analysis of lexical and grammatical learning. Neuroimage. 193, 178–200 (2019).

31. A. D. Friederici, The brain basis of language processing: from structure to function. Physiol. Rev. 91, 1357–1392 (2011).

32. Z. Qi, M. Han, Y. Wang, C. de los Angeles, Q. Liu, K. Garel, E. S. Chen, S. Whitfield-Gabrieli, J. D. E. Gabrieli, T. K. Perrachione, Speech processing and plasticity in the right hemisphere predict variation in adult foreign language learning. Neuroimage. 192, 76–87 (2019).

33. E. Rossi, H. Cheng, J. F. Kroll, M. T. Diaz, S. D. Newman, Changes in white-matter connectivity in late second language learners: Evidence from diffusion tensor imaging. Front. Psychol. 8, 2040 (2017).

34. W. Xin, J. R. Chan, Myelin plasticity: sculpting circuits in learning and memory. Nat. Rev. Neurosci. 21, 682–694 (2020).

35. V. DeLuca, K. Segaert, A. Mazaheri, A. Krott, Understanding bilingual brain function and structure changes? U bet! A unified bilingual experience trajectory model. J. Neurolinguistics 56, 100930 (2020).

36. D. Perani, The neural basis of language talent in bilinguals. Trends Cogn. Sci. 9, 211– 213 (2005).

37. A. D. Friederici, S. M. E. Gierhan, The language network. Curr. Opin. Neurobiol. 23, 250–254 (2013).

38. H. Liu, F. Cao, L1 and L2 processing in the bilingual brain: A meta-analysis of neuroimaging studies. Brain Lang. 159, 60–73 (2016).

39. H. R. P. Park, G. Badzakova-Trajkov, K. E. Waldie, Language lateralisation in late proficient bilinguals: a lexical decision fMRI study. Neuropsychologia. 50, 688–695 (2012).

40. H. Xiang, T. M. van Leeuwen, D. Dediu, L. Roberts, D. G. Norris, P. Hagoort, L2-proficiency-dependent laterality shift in structural connectivity of brain language pathways. Brain Connect. 5, 349–361 (2015).

41. A. D. Friederici, D. Y. von Cramon, S. A. Kotz, Role of the corpus callosum in speech comprehension: interfacing syntax and prosody. Neuron. 53, 135–145 (2007).

42. V. R. Karolis, M. Corbetta, M. Thiebaut de Schotten, The architecture of functional lateralisation and its relationship to callosal connectivity in the human brain. Nat. Commun. 10, 1417 (2019).

43. J. S. Bloom, G. W. Hynd, The role of the corpus callosum in interhemispheric transfer of information: excitation or inhibition? Neuropsychol Rev. 15, 59–71 (2005).

44. L. J. van der Knaap, I. J. M. van der Ham, How does the corpus callosum mediate interhemispheric transfer? A review. Behav. Brain Res. 223, 211–221 (2011).

45. A. Zalesky, A. Fornito, E. T. Bullmore, Network-based statistic: Identifying differences in brain networks. Neuroimage. 53, 1197–1207 (2010).

46. Z. Gracia-Tabuenca, S. Alcauter, NBR: Network-based R-statistics for (unbalanced) longitudinal samples. bioRxiv. 373019 (2020).

47. E. Tschirner, Examining the validity and reliability of the ITT vocabulary size tests. Research Papers in Assessment. Univ. Leipzig. 3 (2021).

48. S. J. Forkel, E. Rogalski, N. Drossinos Sancho, L. D’Anna, P. Luque Laguna, J. Sridhar, F. Dell’Acqua, S. Weintraub, C. Thompson, M. M. Mesulam, M. Catani, Anatomical evidence of an indirect pathway for word repetition. Neurology. 94, e594–e606 (2020).

49. G. Hickok, D. Poeppel, The cortical organization of speech processing. Nat. Rev. Neurosci. 8, 393–402 (2007).

50. J. Legault, A. Grant, S.-Y. Fang, P. Li, A longitudinal investigation of structural brain changes during second language learning. Brain Lang. 197, 104661 (2019).

51. E. F. Lau, C. Phillips, D. Poeppel, A cortical network for semantics: (de)constructing the N400. Nat. Rev. Neurosci. 9, 920–933 (2008).

52. G. Raboyeau, K. Marcotte, D. Adrover-Roig, A. I. Ansaldo, Brain activation and lexical learning: the impact of learning phase and word type. Neuroimage. 49, 2850–2861 (2010).

53. K. Veroude, D. G. Norris, E. Shumskaya, M. Gullberg, P. Indefrey, Functional connectivity between brain regions involved in learning words of a new language. Brain Lang. 113, 21–27 (2010).

54. L. Ghazi Saidi, V. Perlbarg, G. Marrelec, M. Pélégrini-Issac, H. Benali, A. I. Ansaldo, et al., Functional connectivity changes in second language vocabulary learning. Brain Lang. 124, 56–65 (2013).

55. A. M. Grant, S. Y. Fang, P. Li, Second language lexical development and cognitive control: A longitudinal fMRI study. Brain Lang. 144, 35–47 (2015).

56. M. Vigneau, V. Beaucousin, P. Y. Hervé, G. Jobard, L. Petit, F. Crivello, E. Mellet, L. Zago, B. Mazoyer, N. Tzourio-Mazoyer, What is right-hemisphere contribution to phonological, lexico-semantic, and sentence processing? Insights from a meta-analysis. Neuroimage. 54, 577–593 (2011).

57. R. V. Reichle, A. Tremblay, C. Coughlin, Working memory capacity in L2 processing. Probus. 28, 29–55 (2016).

58. Council of Europe, Common European Framework of Reference for Languages: learning, teaching, assessment (Cambridge University Press, 2001).

59. J. C. Raven, J. H. Court, Raven’s progressive matrices and vocabulary scales (Oxford: Oxford pyschologists Press, 1998).

60. B. Fischl, D. H. Salat, E. Busa, M. Albert, M. Dieterich, C. Haselgrove, A. Van Der Kouwe, R. Killiany, D. Kennedy, S. Klaveness, A. Montillo, N. Makris, B. Rosen, A. M. Dale, Whole brain segmentation: Automated labeling of neuroanatomical structures in the human brain. Neuron. 33, 341–355 (2002).

61. M. F. Glasser, T. S. Coalson, E. C. Robinson, C. D. Hacker, J. Harwell, E. Yacoub, K. Ugurbil, J. Andersson, C. F. Beckmann, M. Jenkinson, S. M. Smith, D. C. Van Essen, A multi-modal parcellation of human cerebral cortex. Nature. 536, 171–178 (2016).

62. C. R. Buchanan, M. E. Bastin, S. J. Ritchie, D. C. Liewald, J. W. Madole, E. M. Tucker-Drob, I. J. Deary, S. R. Cox, The effect of network thresholding and weighting on structural brain networks in the UK Biobank. Neuroimage. 211, 116443 (2020).

## SI References

1. Council of Europe, Common European Framework of Reference for Languages: learning, teaching, assessment (Cambridge University Press, 2001).

2. D. Little, The Common European Framework of Reference for Languages: Content, purpose, origin, reception and impact. Language Teaching. 39, 167–190 (2006).

3. E. Tschirner, Aligning frameworks of reference in language testing: The ACTFL proficiency guidelines and the common European framework of reference for languages. (Stauffenburg-Verlag, 2012).

4. E. Tschirner, Examining the validity and reliability of the ITT vocabulary size tests. Research Papers in Assessment. Univ. Leipzig. 3 (2021).

5. N. Weiskopf, J. Suckling, G. Williams, M. M. Correia M., B. Inkster, R. Tait, C. Ooi, E. T. Bullmore T., A. Lutti, Quantitative multi-parameter mapping of R1, PD*, MT, and R2* at 3T: A multi-center validation. Front. Neurosci. 7, 95 (2013).

6. T. E. J. Behrens, H. J. Berg, S. Jbabdi, M. F. S. Rushworth, M. W. Woolrich, Probabilistic diffusion tractography with multiple fibre orientations: What can we gain? Neuroimage. 34, 144–155 (2007).

7. A. M. Dale, B. Fischl, M. I. Sereno, Cortical surface-based analysis: I. Segmentation and surface reconstruction. Neuroimage. 9, 179–194 (1999).

8. M. F. Glasser, T. S. Coalson, E. C. Robinson, C. D. Hacker, J. Harwell, E. Yacoub, K. Ugurbil, J. Andersson, C. F. Beckmann, M. Jenkinson, S. M. Smith, D. C. Van Essen, A multi-modal parcellation of human cerebral cortex. Nature. 536, 171–178 (2016).

9. C. R. Buchanan, M. E. Bastin, S. J. Ritchie, D. C. Liewald, J. W. Madole, E. M. Tucker-Drob, I. J. Deary, S. R. Cox, The effect of network thresholding and weighting on structural brain networks in the UK Biobank. Neuroimage. 211, 116443 (2020).

10. K. H. Maier-Hein, et al., The challenge of mapping the human connectome based on diffusion tractography. Nat. Commun. 8, 1349 (2017).

11. A. Zalesky, A. Fornito, E. T. Bullmore, Network-based statistic: Identifying differences in brain networks. Neuroimage. 53, 1197–1207 (2010).

12. L. García-Pentón, A. Pérez Fernández, Y. Iturria-Medina, M. Gillon-Dowens, M. Carreiras, Anatomical connectivity changes in the bilingual brain. Neuroimage. 84, 495–504 (2014).

